# Operating principles of interconnected feedback loops underlying cell fate decisions

**DOI:** 10.1101/2024.05.24.595855

**Authors:** Abhiram Hegade, Mubasher Rashid

## Abstract

Interconnected feedback loops are prevalent across biological mechanisms including cell fate transitions enabled by epigenetic mechanisms driving phenotypic plasticity of carcinoma cells. However, the operating principles of these networks remain largely unexplored. Here, we identify numerous coupled feedback loops driving phenotypic transition in cancers and CD4+ T cell lineage decisions. These networks have three generic structures, serial type (ST), hub type (HT), and cyclic, which we discover to be the hallmarks of lower- and higher-order dynamics. While networks having ST or cyclic topology exhibit multiple alternative states, those having HT topology enable at most two states. We also show that topologically distinct networks with equal node or loop count exhibit different steady-state distributions, highlighting the crucial influence of network structure on emergent dynamics. Irrespective of the topology, networks with autoregulated genes exhibit multiple states thereby “liberating” network dynamics from absolute topological control. Finally, we identify precise gene interaction targets to restrict the multistable network dynamics to a unique state. Our results thus reveal design principles of coupled feedback loops in enabling multiple alternative states while also identifying perturbations to restrict it. These findings can serve as crucial inputs to comprehend multi-fate decisions of cells and phenotypic plasticity in carcinomas.

## 1. Introduction

Understanding the principles underlying the emergence of diverse and functionally distinct cell types within a living organism is fundamental to unraveling development and disease progression mechanisms [1–3]. As cells undergo proliferation and organize into tissues, their molecular characteristics undergo discrete, step-like changes, leading to the generation of various sequences of unique cell states. These transitions ultimately result in the differentiation of specific functional cell types [4]. Consequently, the development of cells can be conceptualized as the formation of branching cell lineages that yield greater diversity and increasingly specialized cell types [5]. This developmental trajectory though driven by gene interaction networks is also significantly influenced by intercellular signaling among differentiating cells, rendering the process nonautonomous yet self-organizing [6]. At each branch point in a cell lineage, cells face choices between alternative, distinct cell types, with each decision referred to as a *cell fate determination*. The high throughput single-cell transcriptional profiling techniques accompanied by mathematical/computational models of transcription networks have profoundly unveiled some of the complexity in cell fate decisions [7–9]. These models bolster the notion that cell fates can be conceptualized as the stable states of a multistable gene regulatory network (GRN) [10]. Recent advances have highlighted that the topology (structure) of GRNs can substantially dictate the fate decisions of cells and have therefore identified specific hallmarks for multistability (multiple alternative states) [11–13]. For instance, from the network topology perspective, studies have shown that the presence of autoregulated genes [14], positive feedback loops [15–17], and their interconnections [18] are hallmarks of multistability. Further, from the mathematical modeling perspective, some recent studies have reported that multistable systems have a sufficient time delay between gene interactions [19,20] and regulatory functions modeling gene interactions are always convex [21].

One of the simplest networks to exhibit bistability (two phenotypes or steady states) is the toggle switch in which two genes, say X and Y, mutually repress each other to form a double-negative (positive) feedback loop. Under ideal conditions, the toggle switch dynamics converge to one of the two attractors, (*X*^*ON*^, *Y*^*OFF*^) or (*X*^*OFF*^, *Y*^*ON*^) thereby explaining the “either/or” choice between the two cell fates [20]. Practical examples of such circuits are found in both development and diseases. For instance, during hematopoiesis, *GATA*1and *PU*. 1 mutually repress and drive a common myeloid progenitor to either erythroid lineage (*GATA*1^*ON*^, *PU*. 1^*OFF*^) or a myeloid lineage (*GATA*1^*OFF*^, *PU*. 1^*ON*^) [22–24]. Similarly, mutual repression between *ptf*1*a* and *Nkx*6 controls the branch point in the fate map where a pancreatic progenitor decides between two lineages: exocrine (*ptf*1*a*^*ON*^, *Nkx*6^*OFF*^) and endocrine (*ptf*1*a*^*OFF*^, *Nkx*6^*ON*^) [5,25]. On the other hand, a prominent example of cell state transition in diseases is epithelial-mesenchymal transition (EMT)-induced metastasis of carcinoma which is primarily driven by double-negative feedback loops formed between miR200 (micro-RNA) – epithelial marker gene, and Zeb (mRNA) – mesenchymal marker gene, family members [26,27]. The EMT confers non-motile epithelial tumor cells with the characteristics of mesenchymal cells, which are more migratory and invasive. The migrating cancer cells then undergo a reverse mesenchymal-to-epithelial transition (MET) to seed metastatic tumors [27,28]. This circuit explains the “go or grow” mechanism of cancer cells in which “go” phenotype (viz. migratory and invasive) is enabled by (*miR*200^*OFF*^, *Zeb*^*ON*^) and “grow” phenotype (viz. proliferative) is enabled by (*miR*200^*ON*^, *Zeb*^*OFF*^) [29,30].

While positive feedback loops (PFLs) are the cornerstone for binary cell fate lineage decisions, in practice, the robust cellular function and reliable cell decision require diverse cross-talk among the genes, which is manifested through interconnected feedback loops [31,32]. For instance, compared to a single PFL, two coupled PFLs significantly increase the range of cellular conditions in which bistability is observed [5,32]. These interconnected positive feedback loops are widespread in biological systems, including stepwise lineage decision of CD4+ T cells, cell cycle regulation, neuronal cell fate decision, calcium signal transduction, B cell fate specification, and EMT [17,31,33–38]. A thorough understanding of the design principles of the coupled PFLs is therefore crucial to understanding robust cell functioning, cell fate decisions, and diseases such as EMT-induced metastasis of carcinomas.

Here, we identify numerous topologically distinct coupled PFLs intertwined with complex biological networks implicated in EMT-enabled carcinoma phenotypic transition and CD4+ T cell differentiation [11,30,37,39–42]. We define these coupled PFLs as high-dimensional feedback loops (HDFLs), a term coined by combining definitions from Ahrends et al. (2014) and Nordick and Hong (2021) [33,35]. We find that these HDFLs have three types of “global” structures (topology): (i) serial type (ST), in which toggle-switches (positive feedback loops formed by two mutually antagonistic genes) are connected serially like a chain, (ii) hub type (HT), in which several toggle switches accumulate on one common toggle switch to form a hub-type network, and (iii) cyclic, in which end-to-end connected toggle switches form a loop **(Fig. 1)**. We investigated multistability as an emergent property of these networks by analyzing their steady-state (attractor or phenotype) space (see methods). Our findings show that serial and hub-type HDFLs have contrasting dynamics, highlighting a new direction to further our understanding of the mechanism leading to multiple alternative states. The ST HDFLs tend to exhibit multiple alternative states which becomes more pronounced as the network size grows larger. The increase in higher-order stability (here onwards denoted by HoS) is compensated by a decline in the frequency of monostable attractors. In contrast, in hub-type HDFLs, the attractor space is restricted to mono-and bistability and as network size becomes larger, we observe a sharp incline in the frequency of bistable attractors which is compensated by a sharp decline in mono-and HoS. Moreover, irrespective of network topology, autoregulations (self-activated genes) shift the steady state distribution (SSD) towards HoS, which becomes more prominent in ST networks compared to HT networks. This implies that autoregulations lead to a partial loss of the influence of topology on network dynamics.

**Fig 1.**
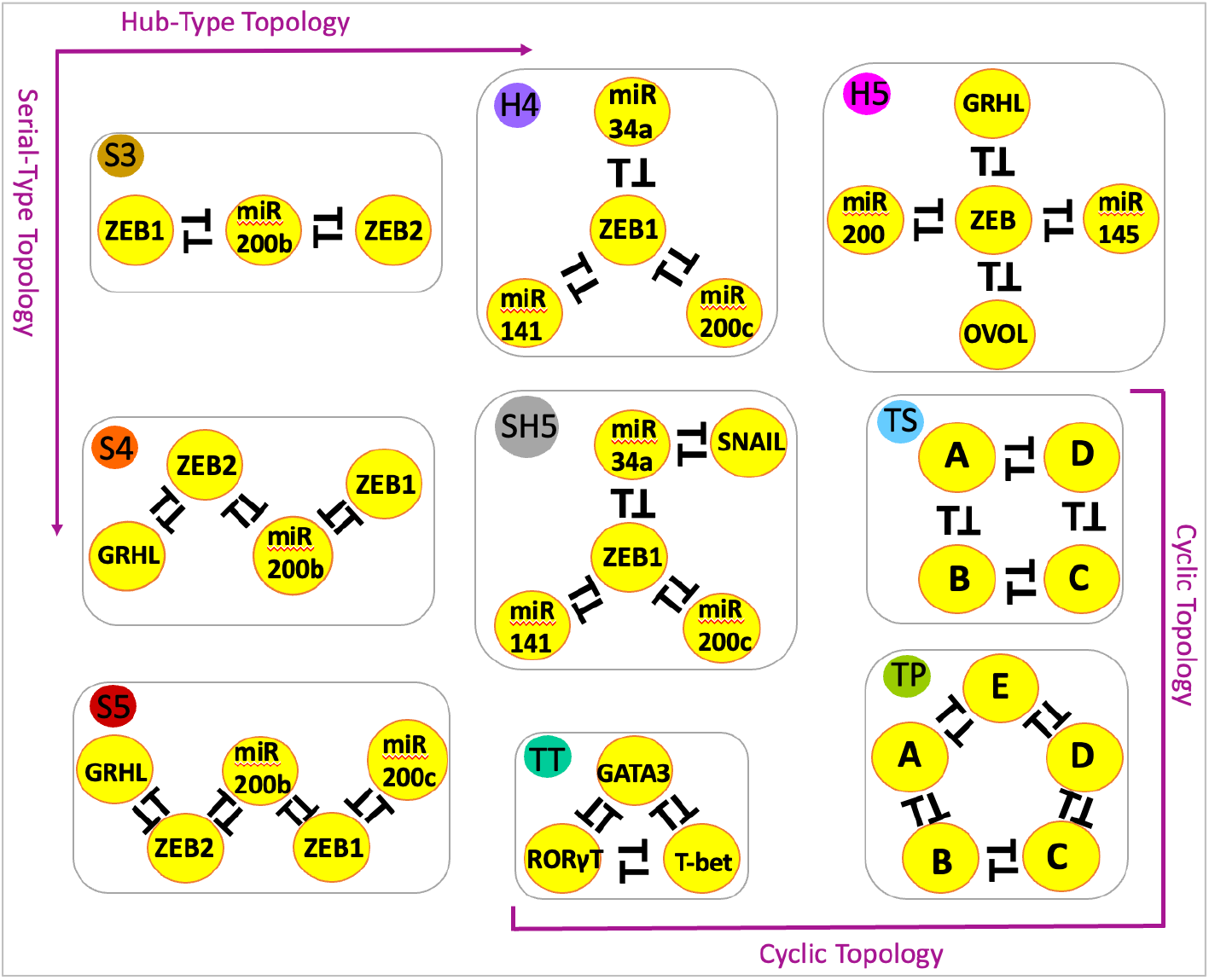
Schematics of high-dimensional feedback loops driving cell fate transition. Networks with annotated nodes are “real” biological networks and those without annotation are “synthetic” networks. In the first column, networks have multiple toggle switches connected end-to-end, forming a serial-type (ST) topology. Along the first row (from S3 to H5 through H4), multiple genes interact with a common node forming a hub-type (HT) topology. In the middle, SH5 has a mixed serial-hub topology. On the bottom right, networks have cyclic topology. Each network is assigned a unique color code for comparative analysis and a name reflecting the type of topology and the number of nodes in the network. For instance, S3, S4, and S5 are ST networks with three, four, and five nodes, respectively. H4 and H5 networks are HT networks with four and five nodes, respectively. SH5 is a mixed serial-hub (SH) network with five nodes. TT, TS, and TP are cyclic networks with three, four, and five nodes, respectively. S3, S4, S5, H4, H5, and SH5 are “real” biological networks driving bidirectional epithelial-mesenchymal fate transitions of carcinoma cells. TT drives cell fate lineage during T-cell differentiation. TS and TP are “synthetic” networks, introduced in the study for comparison purposes. Lines with a bar denote inhibition/repression. Mutual inhibition between nodes is called a toggle switch, which is a PFL.

Next, we constructed all possible topologically distinct HDFLs with three, four, and five nodes that allowed us to introduce “synthetic” networks such as “toggle square” and “toggle pentagon” **(Fig. 1)**. We asked whether HDFLs with the same number of nodes or toggle switches engaged in different topologies can have comparable attractor space. Our analysis reveals that HDFLs with an equal node count or toggle switch count operate differently, highlighting the crucial role of network structure in governing the dynamics. However, we found that cyclic and ST networks with the same node count have comparable dynamics and that cyclic topology further amplifies the frequency of HoS observed in ST networks. To identify targets in the network topology that can restrict the attractor space to lower order, we found that edge sign reversals (converting repressions into activations) that increase negative feedback loops are more effective than edge deletions that reduce PFLs. Finally, we unraveled the distinct effects of these network perturbations in HDFLs with- and without self-activations. While in networks with no self-activated genes, a particular type of perturbation at any position has the same impact, in networks with self-activated genes a perturbation involving the self-activated node significantly increases (reduces) the likelihood of monostable (bistable) states. Together these results highlight the interesting association of network structure with the phenotypic outcome and how autoregulations partially “set free” the dynamics of networks from the topological control. As these networks drive cell fate transitions in development and disease such as metastasis and since multistability can be mapped to phenotypic heterogeneity, our results link the structure of HDFLs with the phenotypic heterogeneity while also suggesting measures to maneuver the structure to control phenotypic plasticity.

## 2 Results

### 2.1 A “tug of war” between serial-and hub-type networks as they pull on the opposite ends of the stability axis

Multiple interconnected toggle switches (referred to as HDFLs throughout this study) often underlie cell decision-making networks, particularly cell fate transition networks governing EMT and MET, and therefore understanding the operational principles of these recurrent network structures can offer promising ways to decode cellular differentiation processes, cell fate switching, and heterogeneity of carcinomas. We curated numerous such networks from the literature and based on their “global” structure, we categorized these into three types: ST (S3, S4, and S5), HT (H4 and H5), and cyclic networks (TT, TS, TP). In ST networks, multiple toggle switches are connected serially or end-to-end to form an extended chain of toggle switches. In contrast, in HT networks, multiple toggle switches are incident on a common central node to form compact networks. With an exception, SH4 has mixed serial/hub type topology **(Fig. 1)**. To understand whether these three network types fundamentally differ in functioning, we used an ordinary differential equation (ODE) based modeling approach implemented in a robust computational algorithm, RAndom CIrcuit Perturbation (RACIPE) [39,43] to analyze the attractor space enabled by each network type **(see Methods)**. Throughout the study, terms like attractors/steady-states/solutions/phenotypes will be used interchangeably.

As depicted in **Fig. 2A**, we observe distinct SSD trends in the ST and HT networks as network size becomes large. The three-node network, S3, allows only one (monostable) and two (bistable) states, the latter being more frequent than the former **(Fig. 2A)**. This shows that the steady-state behavior of two interconnected toggle switches does not differ significantly from that of a single toggle switch, which in ideal conditions also gives rise to at most two steady states. However, we observe that the three interconnected toggle switches formed between four nodes in the S4 network behave differently **(Fig. 2A)**. This network exhibits some tristable (three steady states) attractors, compensated by the decrease in the frequency of monostable attractors, while the frequency of bistable attractors remain nearly unchanged. We asked whether increasing network size by adding more toggle switches serially can further reduce the frequency of monostable solutions and increase the frequency of tristable and other higher-order states. To answer this, we found that in a relatively larger serial network, S5, monostable states decline sharply which is compensated by a sharp increase in tristable states, together with some tetrastable (four states) attractors **(Fig. 2A)**. On the other hand, as we switch to hub-type networks, the SSD of H4 settles around mono-and bistability, where bistable states are more likely to occur than monostable states **(Fig. 2A)**. As network becomes more hub-type, such as the H5 network, we observe a drop (∼10%) in monostable states and a further increase in bistable states while the tri- and higher-order states almost disappear. This begs the question: do serial-type and hub-type networks pull on the lower and higher order ends of the stability axis, respectively? To answer this, we tested our hypothesis on larger (generic) serial and hub networks each having 10 nodes. Strikingly, our findings show that a 10-node serial network has almost equal probability (∼20%) to exhibit bi, tri, and tetrastable states, which is nearly double the probability of monostable states (∼10%). On the other hand, A 10-node hub-type network has almost 85% chance for bistability and 15% chance for monostability **(Fig. S1)**. Our results thus uphold in larger networks and we conclude that ST topology favors HoS that increases (and saturates) in larger networks. In contrast, HT topology favors lower-order stability and bistability becomes dominant in larger hub networks. These findings thus identify design principles of HDFLs exhibiting two and multiple alternate states.

**Fig 2.**
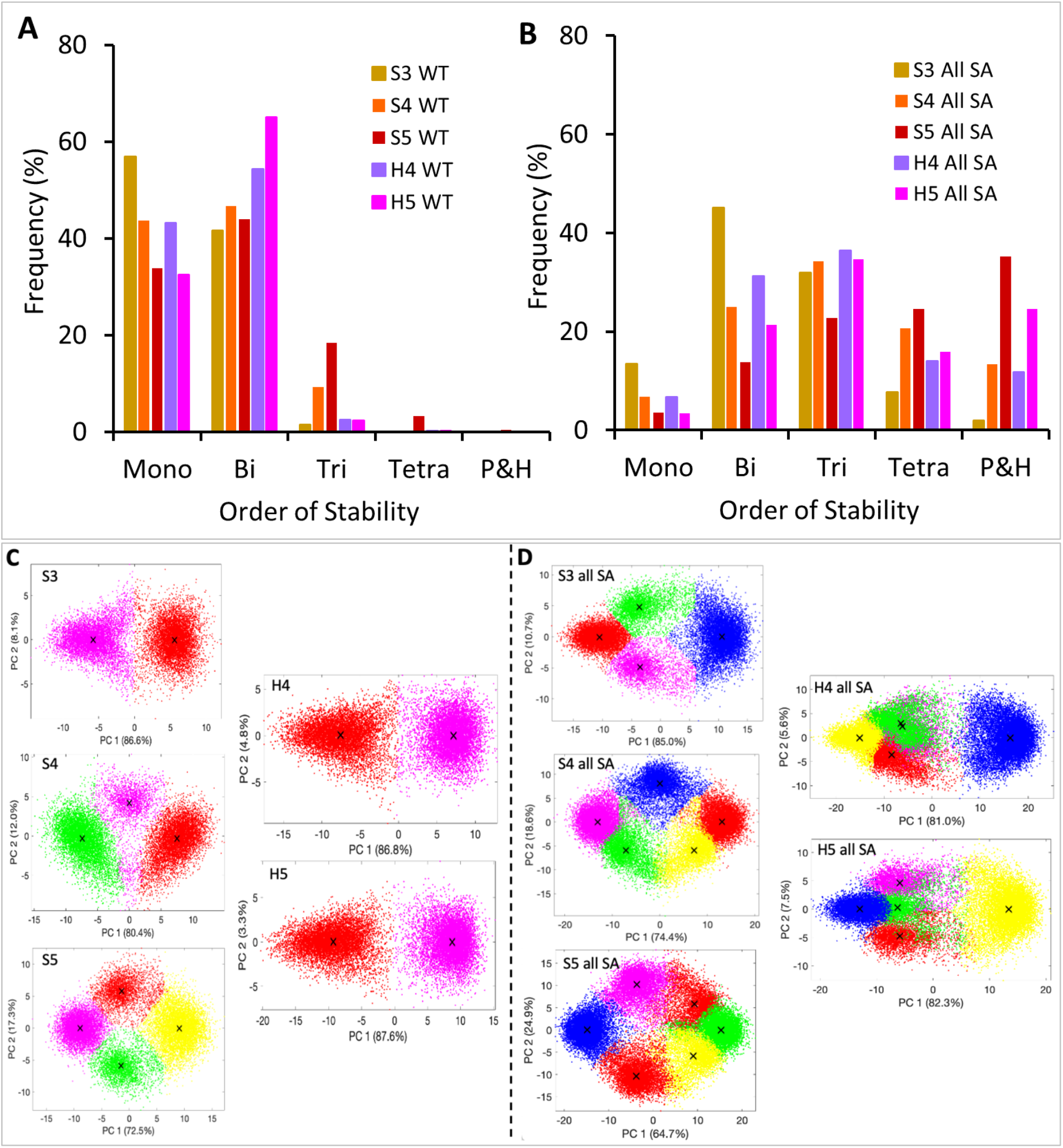
Distribution of stable steady states for serial and hub-type networks. The SSD of wild-type serial-and hub-type networks (A) and their self-activated (SA) counterparts (B). Gene expression data projected on the 1^st^ and 2^nd^ principal component axis for WT networks (C) and their self-activated counterparts (D). Each dot (of 10,000 dots) in the plot is a stable steady state (attractor or phenotype) of a model corresponding to a particular choice of the parameter set. K-means clustering was used to group locally identical attractors and identify distinct “global” attractors that represent distinct cell states (phenotypes) a network can support (see methods for details). The number of phenotypes increase as network size increases serially (say, from S3 to S5), and remains unchanged as network size increases and becomes hub-type (say, from S3 to H4 to H5). Mono, Bi, Tri, and Tetra denote monostable (single), bistable (two), tristable (three), and tetrastable (four) attractors. Penta-and higher (P&H) collectively denotes five and up to ten attractors, respectively and we neglect it throughout the comparative analyses. “WT” implies a wild-type (without any perturbation) network. “All SA” means all network nodes self-activated. “X” in scatter plots marks centroids of clusters. Cluster colors have no correspondence with the colors used to designate networks.

To connect these findings to the state space dimension, we projected the *in silico* gene expression data of these networks on the first two principal components and employed *k-means clustering* to identify the feasible states (clusters) allowed by each of these networks (see methods). We found that as network size increases serially, the number of alternative states (represented by distinct clusters) also increases proportionately **(Fig. 2C)**. However, irrespective of their size, the state space is restricted to just two clusters for HT networks such as H4 and H5 **(Fig. 2C)**. We thus show that numerous alternative states are a consequence of ST topology, which is though restricted by the HT topology. These findings therefore reveal opposing roles of serial-and hub-type topology of networks in controlling alternative fates during cell state transition and highlight that hub-type topology (one manifestation of which can be higher complexity) can, in fact, work against and prevent the occurrence of numerous states. Our findings elucidate how the structural composition of HDFLs can influence the dimensionality of attractor space and concomitant cell fate ‘canalization’ during lineage decision. Interestingly, our observations corroborate with previous studies showing that large networks (which resemble our HT HDFLs), despite their complexity, exhibit only a limited number of viable attractors, a phenomenon termed ‘canalization’ of complex networks [44–47].

We next investigated the impact of self-activations (which are very common in gene regulation) on the dimension of attractor space. To analyze this, we assumed that each gene (node) can activate its transcription and therefore we constructed the self-activated counterparts of these networks. The SSD of networks with all self-activated genes revealed interesting roles of autoregulations in ST and HT networks. As depicted in **Fig. 2B**, for each network, the SSD peak is shifted towards HoS as compared to their WT counterparts **(Fig. 2A)**. A deeper analysis reveals that with the addition of each self-activation, the SSD peak gradually shifts to HoS and a linear relationship between the number of SA genes and the order of stability in each network is evident **(Fig. S2)**. Interestingly, irrespective of the topology, this trend persists across the networks. This finding is further supported by the gene expression data which shows that all self-activated HDFLs exhibit more alternative states (represented by distinct clusters) than their WT counterparts **(Fig. 2D)**. We thus report that autoregulations “liberate” network dynamics from the topological control and evoke a unified response.

### 2.2 Topologically distinct HDFLs with an equal number of nodes operate differently

Of all the “biological” HDFLs reported in **Fig. 1**, we notice that several topologically distinct networks have an equal number of nodes. For instance, S3 and TT each have three nodes, S4 and H4 have four nodes, and SH5 and H5 are five-node networks, integrated through different topologies although. We asked two interesting questions: (i) How many topologically distinct HDFLs can be constructed with three, four, and five nodes, and (ii) Would the SSD of HDFLs with the same number of nodes be similar? We started with three nodes and found that only two distinct HDFLs can be formed: one is the linear chain of three nodes connected with two toggle switches (S3) and the other is a network whose three nodes are the vertices of a triangle connected by three toggle switches, called toggle triad (TT). Likewise, we can only construct three structurally distinct four-node HDFLs, viz. S4, H4, and toggle square (TS), and only four structurally distinct five-node HDFLs, viz. S5, H5, SH5, and toggle polygon (TP) **(Fig. 1)**. This is how “synthetic” networks (in the sense that we don’t have biological examples of these networks) come into play in this study. These cyclic networks (i.e., TT, TS, and TP) have recently been studied in detail by Duddu et al. (2020) [48] and Kishore et al. (2022) [49].

Next, we did a comparative steady-state analysis of networks in each group. When considering three-node networks, we observe that mono-and bistable frequencies in S3 and TT differ slightly and the SSD peak for both networks is skewed towards mono-and bistability. We also notice some tristable attractors (<10%) emerging in TT, which can be attributed to an extra toggle switch in TT **(Fig. 3A1)**. Among the four-node networks, S4 and H4 have comparable mono-and-bistable frequencies but significant differences (∼ 3 times) in tristable frequencies **(Fig. 3B1)**. On the other hand, we find a sharp decline in monostable frequencies in the TS network compensated by an increase in bi-and-tristable frequencies. The tristable frequencies of S3 and TS networks are almost the same. This demonstrates that both cyclic network TS and serial network S4 push for HoS and thus operate differently than the hub network H4. In the five-node networks, it is evident that S5, SH5, and TP have comparable mono, bi, and tristable frequencies, and as expected H5 network has the highest/lowest bistable/tristable frequencies due to its hub-type topology **(Fig. 3C1)**. Also, the SSD of TP shows the emergence of higher tri-and tetrastable solutions compared to other networks in the group. This shows that like TS, the cyclic network TP also operates closer to the “pure” serial network, S5, than hub network H5 or serial-hub network SH5. This difference in dynamics is also reflected in scatter plots characterizing the distinct states allowed by a network for 10,000 different parameter sets **(Fig. S3)**. From the plots, it is evident that among three, four, and five-node networks, cyclic networks exhibit an extra alternative state than serial networks, and the hub networks exhibit just two states **(Fig. S3)**. We thus find that all the topologically distinct networks constructed from a given (same) number of nodes result in a different number of states, typically governed by the specific topology.

**Fig 3.**
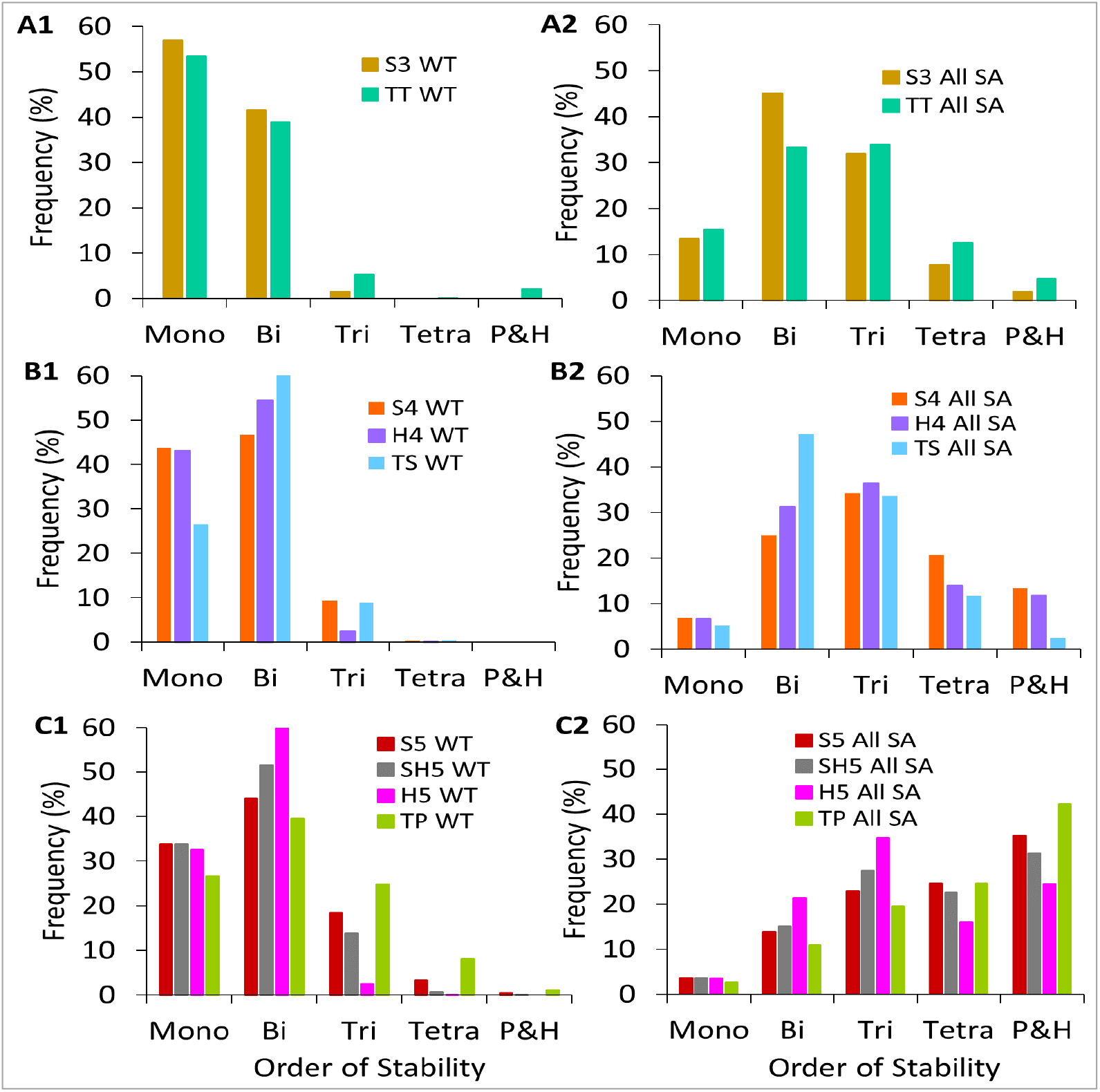
SSD comparison of topologically distinct networks having an equal number of nodes. (A1) Three-node networks showing almost overlapping SSD. (B1) Four-node networks show non-overlapping SSD. Cyclic network TS functions like a serial-type network S3, in the sense that both tend to exhibit higher-order stabilities as compared to H4. (C1) Five-node networks again show non-overlapping bi-and higher-order (tristable and above) SSD. The cyclic network TP functions like a serial network with more frequent higher-order steady states. Except in A1, the same node-count networks in B1 and C1 show variable SSD. (A2, B2, C2) Self-activated counterparts of networks in (A1-C1), show that self-activated networks with the same node count have comparable SSD as compared to their WT counterparts.

We also analyzed the self-activated counterparts of each group of networks, and as expected, irrespective of the network topology **(Fig. 3A2, B2, C2)**, self-activations increase HoS and the SSD moves toward the right as the network nodes increase **(Fig. 3C2)**. Overall, our analysis (for WT) networks demonstrates that, as the network size grows, network topology plays important in shaping the SSD. The serial and hub-type HDFLs with the same node count operate differently. Also, cyclic networks operate more similarly to ST networks than HT networks in a manner that both ST and cyclic topologies favor HoS. In a generic sense, these findings highlight how the same (number of) genes interacting through different topologies can lead to variation in attractor space dimension and exhibit few or multiple alternative cell states.

### 2.3 Topologically distinct HDFLs with an equal number of toggle switches operate differently

In the previous section, we observed that networks with the same number of nodes can have different toggle switch counts. For example, S4 and TS each have four nodes, but TS has one extra toggle switch than S4. We asked whether HDFLs with the same number of toggle switches integrated in different topologies can have comparable SSDs. To answer this, we formed two groups of HDFLs: the first group has three topologically distinct networks, S4, H4, and TT, each having three toggle switches (3TS) and the second group has four topologically distinct networks, S5, SH5, H5, and TS, each having four toggle switches (4TS) **(Fig. 1, Fig. 4)**. Our simulations show that the SSD of 3TS networks is skewed to mono- and bistability. While S4 and H4 have comparable mono-and bistable frequencies, TT has, in contrast, higher monostable and lower bistable frequencies **(Fig. 4A1)**. In the previous section, we noticed that TT operates closer to serial network S3 and each has three nodes, while S4 and H4 each have one extra node than TT. We thus conclude that the difference in SSD of cyclic network TT and serial network S4 is due to the additional node in S4. The same trend is observed in the 4TS networks, where the TS network shows a slight decline in monostable frequencies, compensated by an increase in bi-and-tristable frequencies **(Fig. 4B1)**. On the other hand, as expected, the SSD of self-activated counterparts of these networks is skewed towards the center **(Fig. 4A2)**, in 3TS networks, and towards the right **(Fig. 4B2)**, in 4TS networks, simply due to increased network size by one node. These data show that even HDFLs consisting of the same number of toggle-switches stitched in different topologies can exhibit distinct attractor space. Together with the results in the previous section, we highlight that while having more nodes drives the network dynamics to HoS, however, its is essentially the topology of these interconnected toggle switches that dictates the overall dynamics of HDFLs.

**Fig 4.**
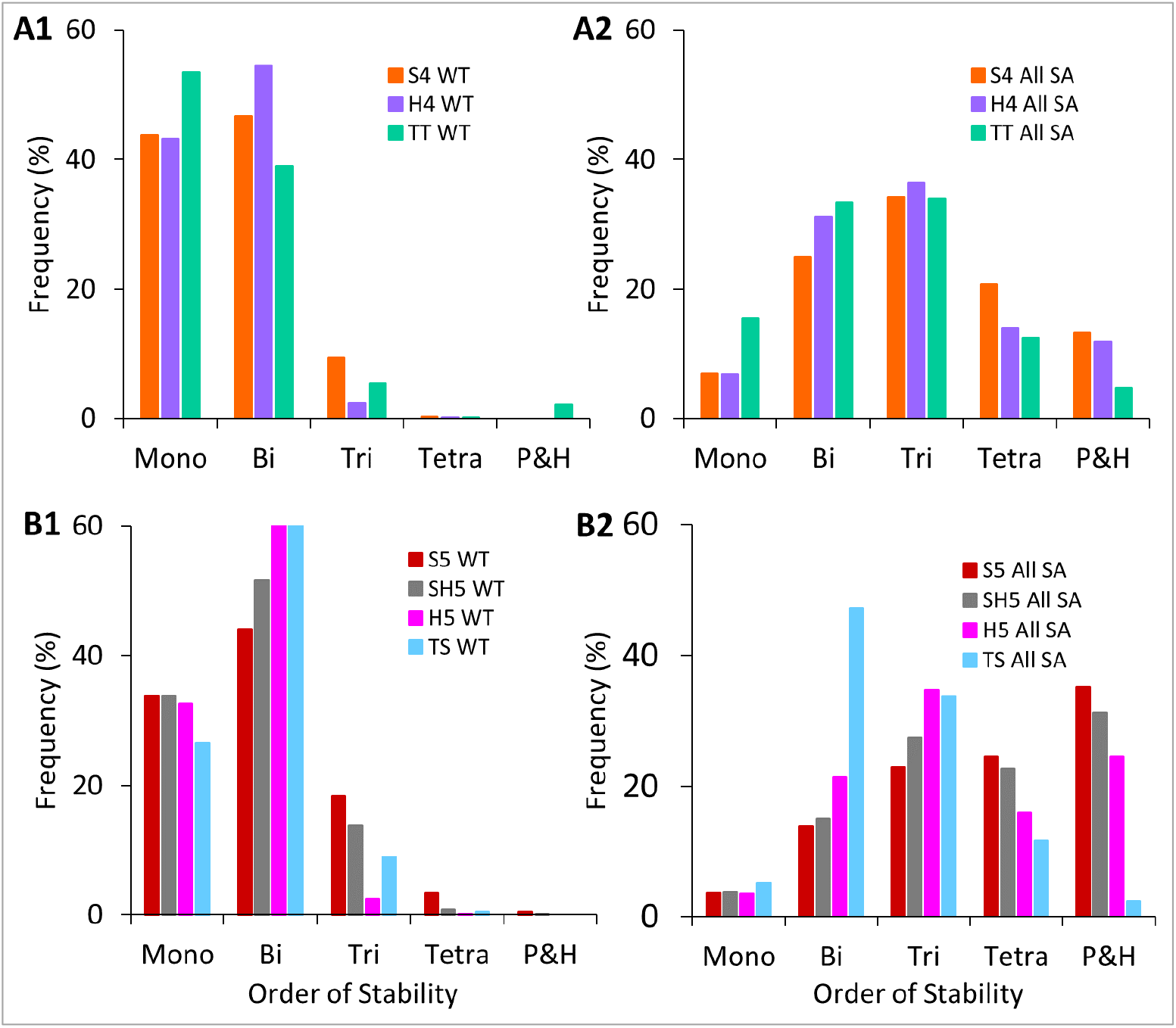
SSD comparison of topologically distinct networks having an equal number of toggle switches. (A1) Networks with three toggle switches have non-overlapping SSD. Despite having one node less than H4, cyclic network TT still shows some possibility for tristability. (B1) Four-node networks again show non-overlapping higher-order SSD. Again TS, despite having one node less than H5, has greater tristable frequencies showing that cyclic network TS functions like a serial-type network by supporting higher-order steady-states.(A2, B2) Self-activated counterparts of networks in (A1) and (B1).

### 2.4 Perturbations to reduce network dynamics to lower-order stability: More negative feedback loops are more effective than less positive feedback loops

As discussed earlier, HDFLs as part of complex networks, drive cell fate transition during differentiation mechanisms and diseases such as EMT-induced non-genetic tumor heterogeneity and carcinoma metastasis. For instance, mutual repression between members of miR200 and ZEB gene families, represented by the S3 network, can drive a carcinoma cell to at least three phenotypes: mesenchymal fate (low miR200, high ZEB), hybrid epithelial-mesenchymal fate (medium miR200, medium ZEB), epithelial fate (high miR200, low ZEB) [30]. Similarly, mutual repression between three genes, RORγT, GATA3 and T-bet, depicted by TT, can drive naïve CD4+ T cell into three distinct states—Th1 (high T-bet, low GATA3, low RORγT), Th2 (low T-bet, high GATA3, low RORγT) Th17 (low T-bet, low GATA3, high RORγT) [48]. The fundamental component that recurs in all of these HDFLs is the PFL formed by two mutually antagonistic genes also called a toggle-switch **(Fig. 1)**. And, so far, we know that more toggle-switches connected serially (like in S5), result in numerous alternative states. We asked, can we target these PFLs to restrict the attractor space of these networks to a few attractors To identify such targets (gene interactions), we introduce two types of edge perturbations **(Fig. 5)** across the networks. The first is the “edge-deletion” (ED) perturbation, where we delete an edge, one at a time, between the nodes and repeat the process as long as the network remains intact. Each ED perturbation thus gradually reduces the number of PFLs in the network. In the “edge-sign-reversal” (ESR) perturbation, we change the sign of one of the inhibitory interactions between the two nodes to activation. Thus unlike ED, each ESR perturbation converts a PFL into a negative feedback loop (NFL) so that more ESRs imply more negative feedback loops **(Fig. 5)**. These two perturbations applied on any network with N toggle switches will give rise to 2N perturbed networks. The implications of these edge perturbations have been tested *in vivo*, for instance, in the case of an EMT PFL involving mutual inhibition of miR200 and Zeb genes. Disrupting specific *miR200c* sites of the endogeneous *Zeb1* locus prevented repression of *Zeb*1 by *miR200c* that resulted into an altogether different functional response [50].

**Fig 5.**
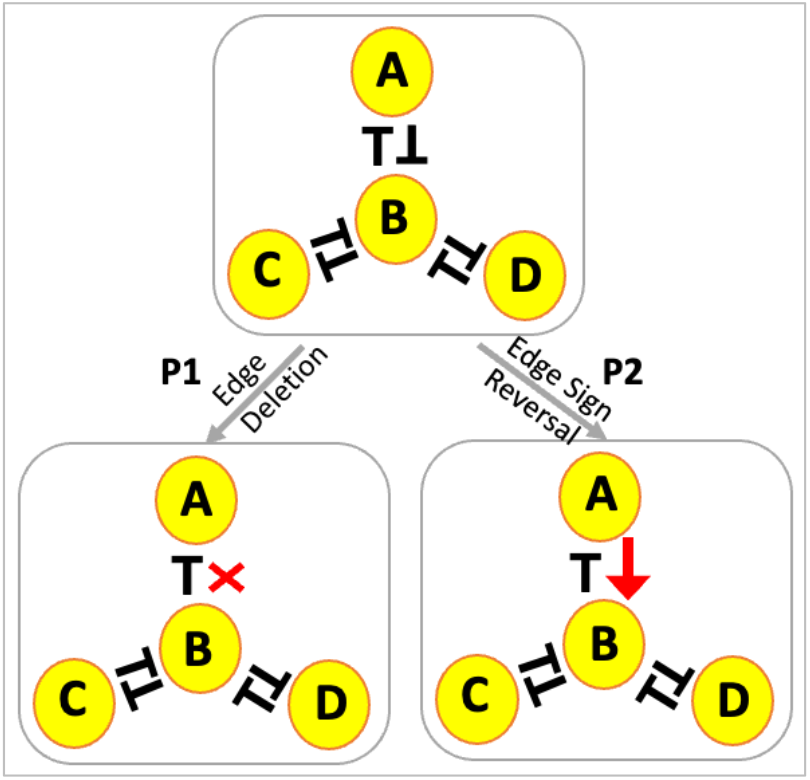
Illustration of the two types of network perturbations. We take any unperturbed network (also called wild type (WT)) and do types of perturbations: Edge deletion (P1) and Edge sign reversal (P2). In P1, one of the edges between the two nodes is deleted, resulting in a breakdown of the positive feedback loop. In P2, the sign of one of the edges between the two nodes is changed from inhibition to activation so that a positive feedback loop transforms into a negative feedback loop. The lines with a bar denote inhibition/repression and those with an arrow denote activation. “X” denotes the deletion of an edge.

We simulated these networks like before using RACIPE and compared the SSD of each perturbed network with its WT (unperturbed) counterpart. Our findings show that both ED and ESR perturbations reduce network SSD to lower-order stability. Particularly, a single ED or ESR increases the frequency of monostable attractors which is compensated by a decrease in the frequency of HoS **(Fig. 6)**. This trend follows throughout the networks and we observe that an ESR is more effective than ED, in the sense that each ESR increases more monostable solutions (by percentage) than an ED **(Fig. S4 A1-C1, Fig. S5 D1-E1, Fig. S6 F1-H1)**. On the other hand, we observe that when all the genes (network nodes) are autoregulated, the same two perturbations have less effect on the SSD of the network **(Fig. S4 A2-C2, Fig. S5 D2-E2, Fig. S6 F2-H2)**. This is evident from the expression data which shows that the number of feasible states (clusters) decrease with an ED or ESR in WT networks (**Fig. 6C)**, which however remain unchanged in their self-activated counterparts **(Fig. 6D)**. We thus conclude that ESR (which increases an NFL) and ED (which decreases a PFL) perturbations work only in non-autoregulated networks to reduce the network attractor space of the network. These edge perturbations can serve as inputs to future efforts focussing on in-depth understanding and controlling multiple cell lineages in differentiation and importantly in preventing phenotypic heterogeneity enabled during the EMT-induced metastasis of carcinomas.

**Fig 6.**
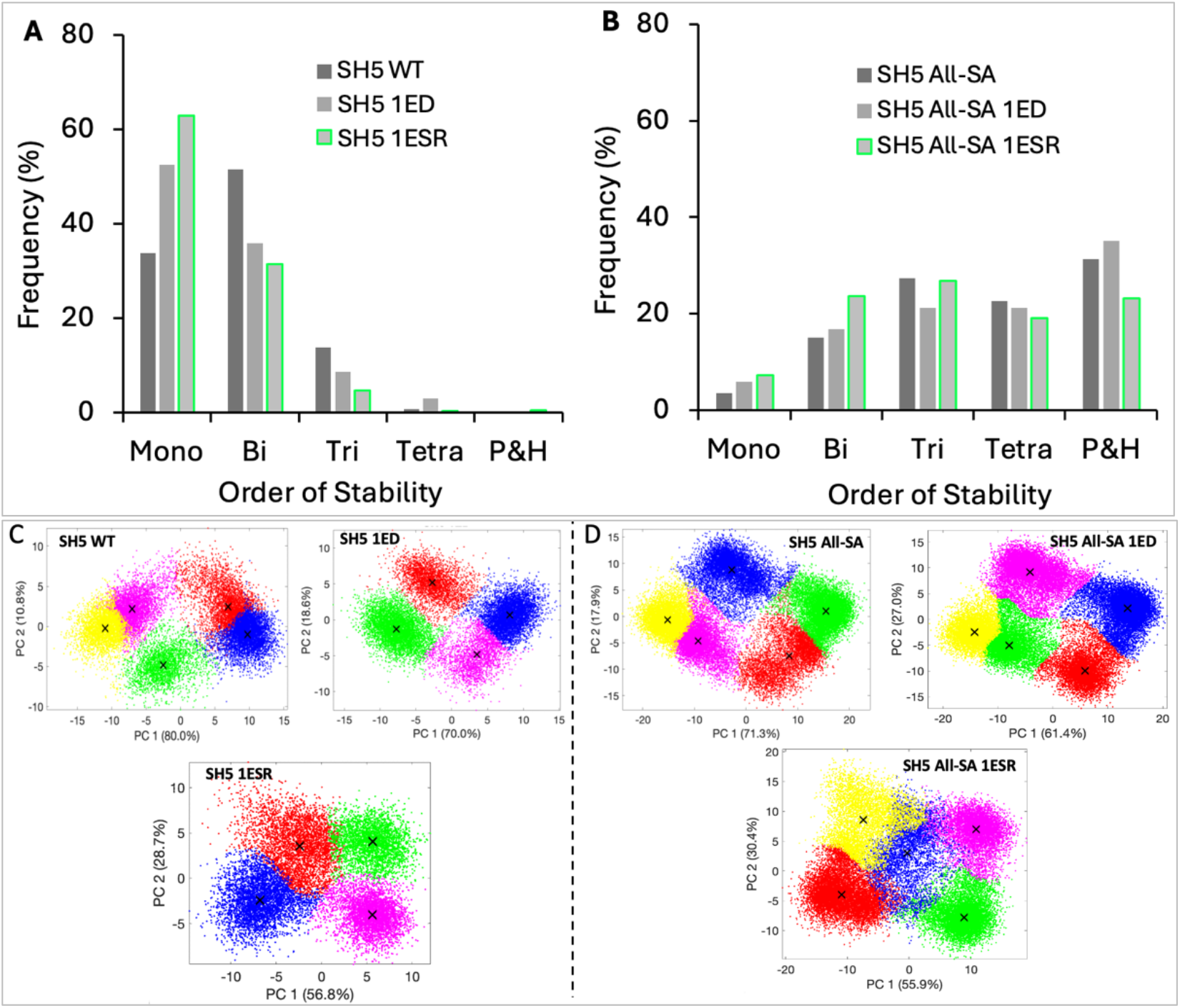
Effect of two types of edge perturbations, ED and ESR, on the SSD of SH5 (representative network). (A) In a non-autoregulated (without self-activations) network, an ESR increases monostable attractors more rapidly than an ED. (B) The effect of edge perturbations is diluted by the self-activations as the differences in SSD between WT and its two perturbed counterparts are negligible. Gene expression data of WT (C) and SA (D) networks projected on the 1^st^ and 2^nd^ principal component axis after clustering. Each cluster represents a network state (or attractor). SH5 WT means SH5 unperturbed network, 1ED means one edge detection, 1ESR means one edge sign reversal, and all-SA means all nodes self-activated.

### 2.5 Identifying perturbations to convert multi-attractor networks to a unique attractor networks

In the previous section, we found that an ED or ESR increases the likelihood of lower-order stability. Taking, for instance, SH5 as a representative network, we show that for the SH5 (wild type) network bistable attractors are the most frequent (∼50%), followed by monostable (∼35%) and tristable (∼15%) attractors. This implies that SH5 likely enables two cell states and in rare cases, three states **(Fig. 7A)**. A single ED in SH5 results in a sharp increase in the frequency of monostable attractors (second bar in Fig. 7A) compensated by a decrease in bi-and tristable attractors. We wanted to analyze the effect of increasing edge perturbations on the dimension of attractor space and asked how many perturbations can turn the network dynamics into a single attractor. To analyze that we gradually increased edge perturbations in the network and noted the SSD after each additional perturbation. We find that as EDs increase, the frequency of monostable attractors also increases linearly, while the frequency of bi-and tristable attractors decrease sharply **(Fig. 7A)**. We also discover that three and four EDs, respectively, eliminate the chance of tristability and bistability, thereby turning SH5 a monostable (single attractor) network. This is also evident in the scatter plot of 10,000 steady states all of which essentially represent a single attractor **(Fig. 7D)**. Since each ED eliminates a PFL and SH5 network has four PFLs, our data thus shows that the removal of all PFLs turns SH5 into a single attractor (monostable) network.

**Fig 7.**
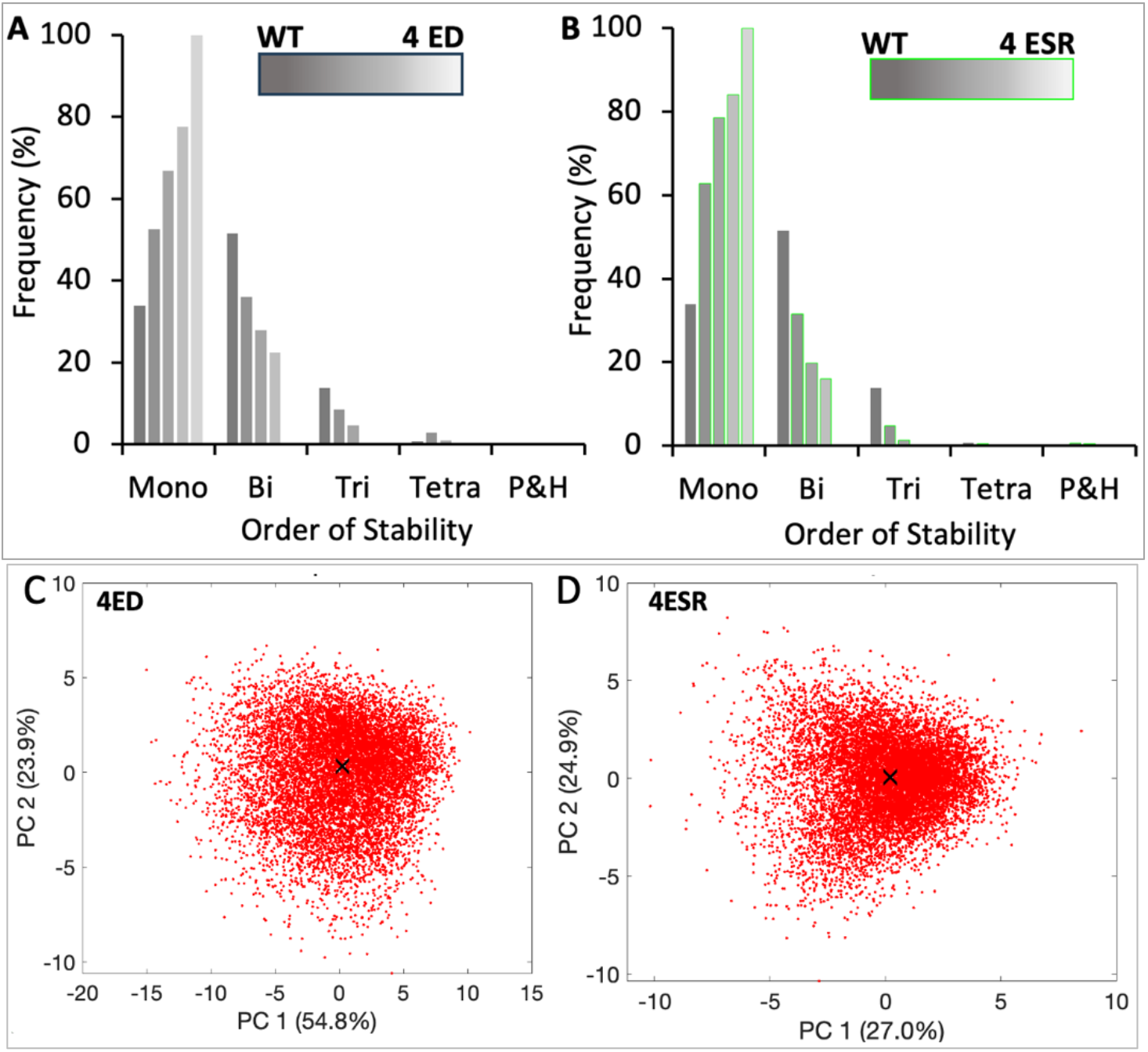
Effect of multiple ED and ESR perturbations on multistability. The plots illustrate the effect of increasing EDs and ESRs on the emergent SSD of HDFLs by considering SH5 as a representative network. (A) As EDs increase, the frequency of monostable attractors increases while bi-and-tri stable attractors decrease proportionately. Deleting three edges (breaking three PFLs) increases the frequency of monostable attractors to nearly 80% from nearly 30% in WT while also removing the chance of occurrence of any tristable solution. A completely monostable dynamics is achieved upon breaking all four PFLs in SH5. (B) As ESRs increase, network dynamics shift to mono-and-bistability faster than in the case of each ED. The 80:20 ratio of monostable: bistable attractors is achieved with two ESRs (i.e., converting two PFLs to NFLs) compared to three EDs and the frequency of tristable attractors is almost negligible. A completely monostable dynamics is achieved by converting four PFLs to NFLs. Gene expression data of SH5 with 4EDs (C) and 4ESR (D) projected on the 1^st^ and 2^nd^ principal component axis after clustering. A cluster represents a single attractor state. WT denotes no ED or ESR and 4ED (4ESR) denotes four edge deletions (edge sign reversals).

Next, we find that increasing ESRs also drive monostable attractors while reducing the frequency of HoS **(Fig. 7B)**. However, compared to EDs, the ESRs induce a rapid decline in bi-and tristable attractors and a surge in monostable attractors compared to ED. This is further supported by the fact that just two ESRs (compared to three EDs) are required to eliminate tristable attractors in SH5 **(Fig. 7B)**. However, four ESRs are required to attain a unique attractor (fifth bar in Fig. 7B shows 100% monostability) which also reflects from the scatter plot of the expression data in **Fig. 7D**. The single cluster observed in the scatter plot shows that all the 10,000 steady states in SH5 network with 4ESR essentially represent a single attractor state. Our findings thus reveal that breaking all the (positive) feedback loops in a network results in a single attractor dynamics. This data is consistent with the traditional studies relating positive feedback loops with multistability.

The positive (negative) correlation between multistability and PFLs (NFLs) is observed across the HDFLs **(Fig. S7 A1-H1)**. However, in their self-activated counterparts, perturbations shift the SSD peak to the left (lower-order stability), and with maximum perturbations, the peak settles around bi- and tristability **(Fig. S7 A2-H2)**. This isn’t surprising, since we have previously shown this correlation of multistability with PFLs and NFLs in large and complex EMT networks as well [11]. We thus conclude that regardless of the network size and topology, PFLs and NFLs have opposite effects on multistability: more PFLs favor HoS while more NFLs enable lower-order stability. Our results therefore provide interesting insights into the association of topology with emergent multistability and can have clinical implications to decode the mechanism enabling multi-fate cell lineages and heterogeneity of carcinomas.

### 2.6 Position of edge perturbation has significance in networks with self-activated nodes

The HDFLs with autoregulated (self-activated) genes are more likely to exhibit multiple alternative states than those without autoregulations. One can then expect that the same perturbation can have a differential effect in auto- and non-autoregulated networks. To investigate this, we first tested our hypothesis on non-autoregulated HDFLs by considering S3 as a representative case. In this network, we deleted an edge in each toggle switch (referred to as P1 and P2 positions), one at a time. We simulated the two perturbed networks and compared the SSD with their WT counterpart. Our data shows that both EDs result in similar SSD, implying that an ED at any position in a network will have the same effect **(Fig. 8A)**. Next, we did the two ESRs in two toggle switches, one at a time. Strikingly, our data shows a complete overlap of two SSDs **(Fig. 8B)**. We thus conclude that, in HDFLs without autoregulated genes, the same type of edge perturbation at two different positions creates a same impact and therefore the position of perturbation has no significance.

**Fig 8.**
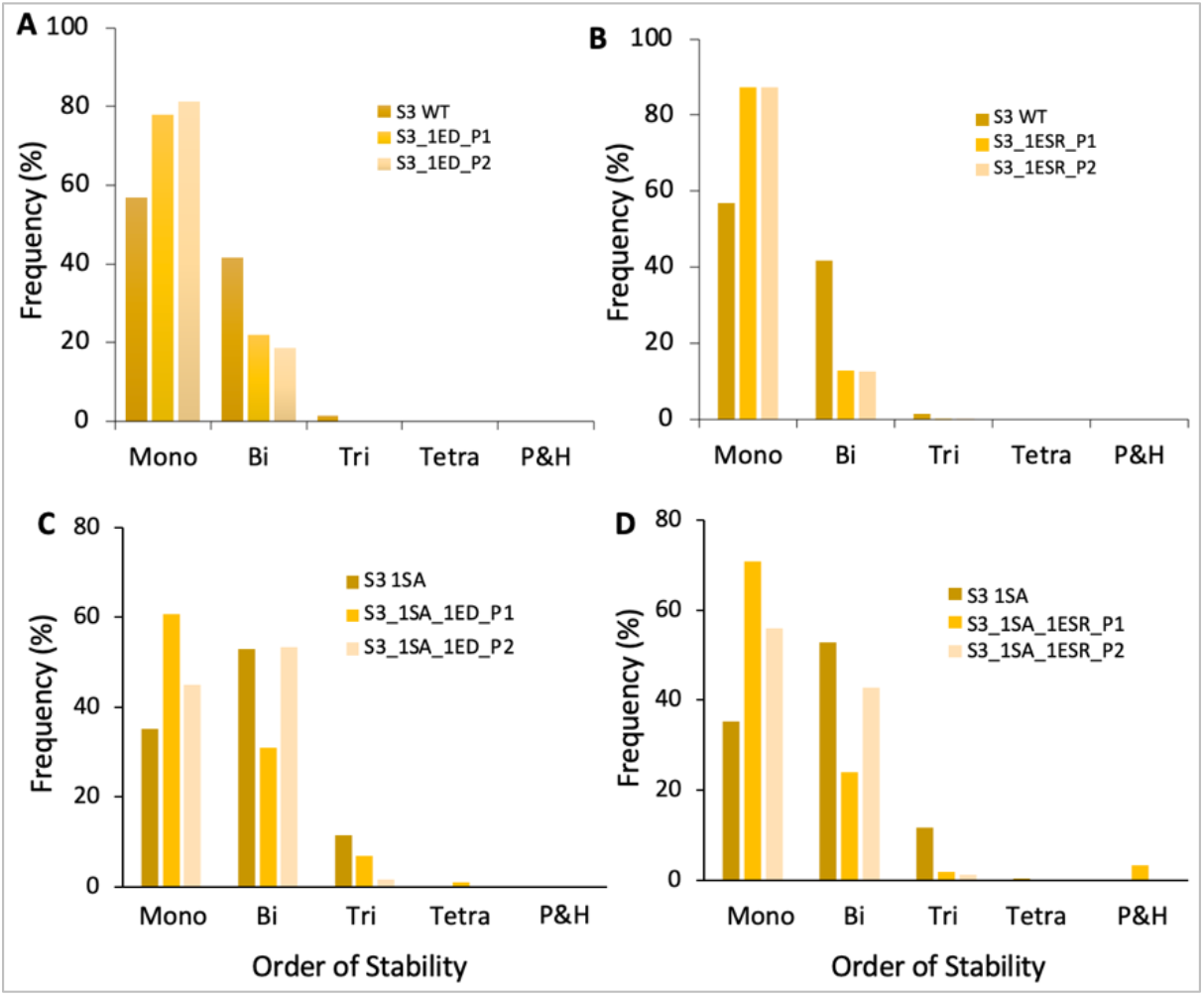
Bar plots showing the importance of the position of edge perturbations in networks with- and without self-activated nodes. (A) A non-self-activated (WT) S3 network and its two ED perturbations. (B) Same as (A) but with ESRs. (C) S3 network with one of the terminal nodes self-activated and its two ED perturbations. (D) same as (C) but with ESRs. Perturbations, P1 and P2, are done in two different toggle switches of S3, so that in (C) and (D), P1 involves the SA node while P2 doesn’t. 1EDP1/1EDP2 (1ESRP1/1ESRP2) denote single-edge deletions (edge sign reversals) at P1 and P2 position. This terminology follows throughout the bar plots in Fig. S8.

To test our hypothesis on HDFLs with autoregulated nodes, we self-activated one of the terminal nodes (ZEB1, in this case) of S3. With SA node, the two PFLs on either side of the central node become “structurally” different - one involves a SA node and the other doesn’t. We found that perturbing an edge (ED or ESR) in the PFL involving the self-activated node (denoted by P1 position) significantly increases (decreases) the likelihood of monostable (bistable) states than an edge perturbation in PFL without self-activated nodes (P2 position). As shown, EDs at P1 position increase nearly 15% monostable states and decrease nearly 20% bistable cases than an ED at position P2 **(Fig. 8C)**. This effect persists when EDs are replaced by ESRs (**Fig. 8D)** and is also observed across HDFLs **(Fig. S8)**. Our findings thus highlight that in networks involving self-activated genes, disrupting the binding interactions between self-activated genes can be more effective in restricting the network dynamics to fewer alternative states (attractors) than disrupting the interactions between non-self-activated genes. These findings thus reveal crucial targets to, for instance, control cell fate transition in diseases such as restricting phenotypic heterogeneity in carcinomas.

## 3. Discussion

Different network structures with specific dynamical behavior underlie a variety of cell fate decision-making programs including epithelial mesenchymal plasticity. Among these, positive and negative feedback loops are ubiquitous and widely studied network motifs with very well-known functionality. For instance, canonical positive feedback loops are responsible for hysteric behavior while negative feedback loops display oscillatory response. We identified numerous HDFLs intertwined in complex GRNs driving EMT-induced cell fate transition during metastasis and CD4+ T cell lineage decision. These HDFLs, composed of interconnected PFLs, have three different “global” topologies: Serial, Hub, and Cyclic. Serial-type networks are formed of serially connected positive feedback, in hub-type networks, many positive feedback loops are incident on a common node forming a relatively dense network structure, and cyclic networks form a ring by a connection between end nodes. We then investigated the emergent dynamics of these networks by particularly looking at the distribution of the possible attractor space. Our analysis revealed contrasting operating principles of ST and HT networks that is predominantly driven by network topology. While ST networks favor HoS, HT networks favor lower-order stability. To investigate the significance of self-activations in each network topology, we found that self-activations somehow set the network dynamics free from topological control and that all self-activated ST, HT, and cyclic networks support HoS.

We further analyzed the significance of node and loop count on the emergent dynamics of HDFLs. We asked whether all possible networks with an equal number of nodes or toggle switches but distinct topologies can have similar operating principles. To answer this we constructed all topologically distinct HDFLs with three, four, and five nodes for comparison. We found that networks with the same node count as well as those with the same loop count have significantly different SSDs and the difference further increases as the network size grows. We also found that cyclic networks operate closer to ST networks compared to HT networks. This analysis thus revealed general operating principles of networks with distinct topologies and upholds the crucial role of topology in the overall emergent dynamics of HDFLs.

Next, we were interested in identifying crucial targets to reduce higher-order network dynamics to lower-order dynamics. We, for instance, asked, what would it require to restrict network dynamics to monostability or bistability. To answer this, we perturbed networks with EDs (breaking PFLs) and ESRs (introducing NFLs) and compared the SSD of perturbed networks with their “WT” counterparts. While EDs and ESRs both reduce the network dynamics to lower-order stability, the effect of ESRs is more rapid and pronounced than EDs. This shows that breaking PFLs has different dynamical implications than converting PFLs to NFLs and that more NFLs in a network likely results in lower-order stability.

Finally, we investigated the significance of the position of the perturbation in HDFLs with non-self-activated and self-activated genese. Our analysis revealed that the position of edge perturbation doesn’t have any major significance in networks with no self-activated genes. However, strikingly, in networks with self-activated genes, an edge perturbation in the loop that involves a self-activated node is more effective in reducing network dynamics to lower-order stability than an edge perturbation in the loop involving two non-self-activated nodes. This effect is more pronounced in ESRs compared to EDs. Collectively, our findings unveil the operating principles of topologically distinct HDFLs by shedding light on the intricacies of topology-dynamics associations. These insights into how network topology, size, and position of perturbations differentially influence emergent dynamics. Our findings can offer an initial guess about the dynamics of complex networks involving HDFLs without without actually simulating their dynamics. It also paves the way to precisely manipulate network properties such as PFLs and NFLs in complex networks to control dynamical characteristics particularly the order of stability as discussed in this work. Our study offers valuable insights into the crucial role of network topologies in driving the emergent dynamics of HDFLs while also identifying critical inhibitors and their positions with respect to the topology to restrict the higher-order dynamics. Since many of these networks underlie cell fate programs in development, differentiation, and diseases like EMT-enabled carcinogenesis, our results thus have implications in comprehending, for example, cell lineage switching checkpoints, besides identifying inhibitors to curtail EMT-enabled phenotypic heterogeneity and metastasis of carcinoma.

## 4. Methods

### 4.1 Numerical simulations

The numerical simulations are performed in RAndom CIrcuit PErturbation (RACIPE) https://github.com/simonhb1990/RACIPE-1.0 which is a robust computational method to simulate the dynamics of complex transcriptional networks [51]. The input to RACIPE is the network topology file containing descriptions of genes (nodes) and their interactions (edges) which is converted into a system of coupled ODEs. The number of variables in the mathematical model corresponds to the number of genes in the network. For instance, given a network with *A*_*i*_ activating nodes and *R*_*j*_ inhibiting nodes incident on a node *X*, the dynamics of *X* is modeled using the following nonlinear ordinary differential

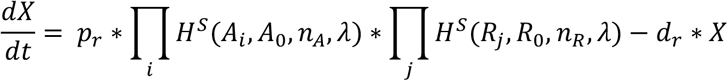

Where *X, A*_*i*_, and *R*_*j*_ represent concentrations, and *p*_*r*_ and *d*_*r*_ the maximum production and degradation rates. The *H*^*S*^ is the shifted Hill function defined as,

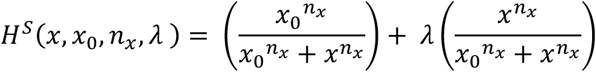

Where *x*_0_ represents the activation/inhibition threshold, *n*_*x*_ the Hill coefficient, and *λ* the fold change in activation/inhibition. The generic form of the shifted Hill function, *H*^*S*^, provides a unified, single-function description for both activators and inhibitors for different ranges of *λ*.

We define,

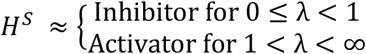

The kinetic parameters associated with the mathematical model are randomly sampled from reasonable predefined ranges listed in **Table 1** using **log-uniform distribution**. The mathematical model corresponding to each network is solved numerically using the ***Euler*** method of integration with a step size of ***0*.*1, 100*** initial conditions, and ***20*** iterations for each initial condition. The cut-off for convergence of steady state (time-invariant solution) is fixed at ***1*.*0***. Simulation of all the networks is done in triplicates, with each replicate having **10000** parameter sets, which takes into account cell-to-cell heterogeneity in biochemical reaction rates. For each network, the output is a data set in which columns correspond to genes and the rows correspond to the stable steady states. This data takes a form similar to typical experimental *microarray* data and we thus employ clustering techniques (discussed below) to identify the total number of feasible states allowed by a network.

**Table 1.** Ranges of parameters that are random sampled using log-uniform distribution. [.] represents closed interval and (.) represents open interval.

| Parameter | Range |
| --- | --- |
| Production Rate ($p_r$) | [1, 100] |
| Degradation Rate ($d_r$) | [0.1, 1] |
| Fold change of Activation ($\lambda$) | (1, 100] |
| Fold change of Inhibition ($\lambda$) | [0.01, 1) |
| Hill Coefficient ($n_x$) | [1, 6] |
| Threshold value ($x_0$) | The threshold value is chosen so that the node has a 50% probability of being active (half-functional rule) |

### 4.1 Data clustering and projection

The in silico gene expression obtained from numerical (RACIPE) simulations is normalized using Z-score standardization method given by the following equation.

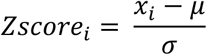

Where the mean (*μ*) and the standard deviation (*σ*) is taken over all the 10,000 models.

#### Principal Component Analysis (PCA)

The PCA was used to project the data on the two principal axis explaining maximum percentage of the total data variance. We used singular value decomposition to perform PCA.

#### K-means Clustering

We used K-means clustering, an unsupervised learning algorithm, with *L*^1^ ***norm*** (defined below) to segregate the data into different clusters and identify the feasible alternate states.

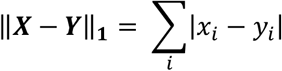

for *X* = (*x*_1_, *x*_2_, *x*_3_, …, *x*_*n*_) and *Y* = (*y*_1_, *y*_2_, *y*_3_, …, *y*_*n*_)

To make sure that the clusters identified by the algorithm were correct, the algorithm was run **1000** times for each data set. The best clustering outcome was chosen to be the one with a minimum value of *“Within Cluster Sum of Squares (WCSS)”*.

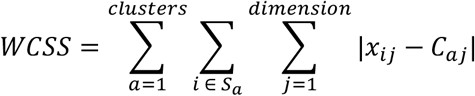

Where the first two summations calculate distances of data points in a cluster from the centroid, C, of that cluster and the third summation takes the sum over all clusters.

## Author contributions

**A.H**.: Simulations, data analysis, discussions. **M.R**.: Conceptualization, supervision, simulations, data analysis, discussions, data presentation, writing, funding acquisition.

## Data and Code Availability

All the raw data generated for this study and the codes for the analysis of RACIPE data are available on the GitHub page: https://github.com/AbhiramHegade/

## Acknowledgments

M.R. gratefully acknowledges funding from the following agencies of Govt. of India.

(1) Department of Science and Technology (DST), Grant No. DST/INSPIRE/04/2020/001492
(2) Science and Engineering Research Board (SERB), Grant No. CRG/2023/006432

