## Supplementary Material for "Operating principles of interconnected feedback loops underlying cell fate decisions"

**S1. Topology-dependent steady-state distribution in large serial and hub-type networks**

In this section, we constructed larger size (10 nodes) ST and HT HDFLs and analyzed the probability distribution (in percentage) of mono, bi, tri, and tetrastable states. We show that our findings for small ST and HT HDFLs with three, four, and five nodes also hold for larger HDFLs. As shown in Fig. S1A, the serial network exhibits equal (~20%) possibilities for bi, tri, and tetrastable states and quite a lesser chance (~10%) for monostability, implying that serial-type topology favors higher-order stability. In Fig. S1B, however, we see that the hub-type topology leads to bistability (~85%) with a little chance for monostability (~15%). This indicates that larger hub-type networks are strictly bistable systems.

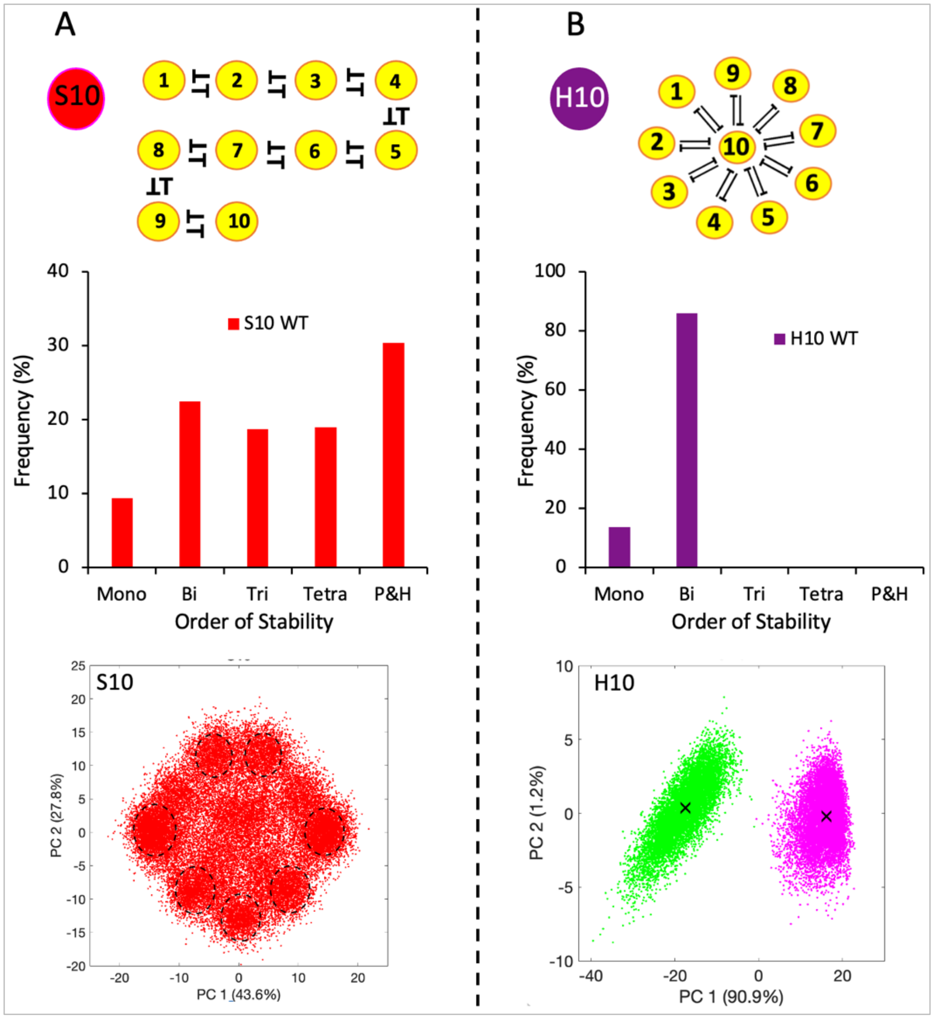


*Fig. S1 Schematics and SSD of larger serial and hub type HDFLs, each having ten nodes and nine toggle switches. (A) The ST topology exhibits almost equal (~20%) possibility for bi, tri, and tetrastable states with the occurrence of some (<10%) monostable states as well. (B) The HT topology shows complete dominance of bistability (~85%) over monostability (~15%) while also eliminating the chance for tri-and tetrastable states. P&H denotes penta-and higher-order states that collectively include five and up to ten states. The lower horizontal panel shows in silico gene expression projected on the 1$^{st}$ and 2$^{nd}$ principal component axes and clustering of multiple states in S10 and just two states in H10.*

**S2. Autoregulations enable higher-order stability.**

In this section, we investigate the effect of autoregulations on network dynamics. After self-activating a particular node in a network, we simulate its emergent dynamics and compare the steady-state distribution with the corresponding WT (no self-activated nodes) network. We repeat this process until all the nodes in the network are self-activated. In the plots below, the first bars (dark color) corresponding to each order of stability, i.e., mono, bi, tri, tetra, and P&H stability, compose the SSD of the WT network. The bars at the second place compose the SSD of the network with a single node self-activated, the bars at the third place compose the SSD of the network with two nodes self-activated, and so on. As shown, as more nodes get self-activated, the frequency of mono-and bistable attractors decrease which is compensated by increase in the frequency of tri-and tetrastable attractors. The peak of SSD thus shifts from lower-order to higher-order stability. Interestingly, this trend becomes more evident as network size increases and can be noticed throughout the networks. We thus conclude that regardless of the topology, networks with self-activations tend to exhibit multiple alternative states.

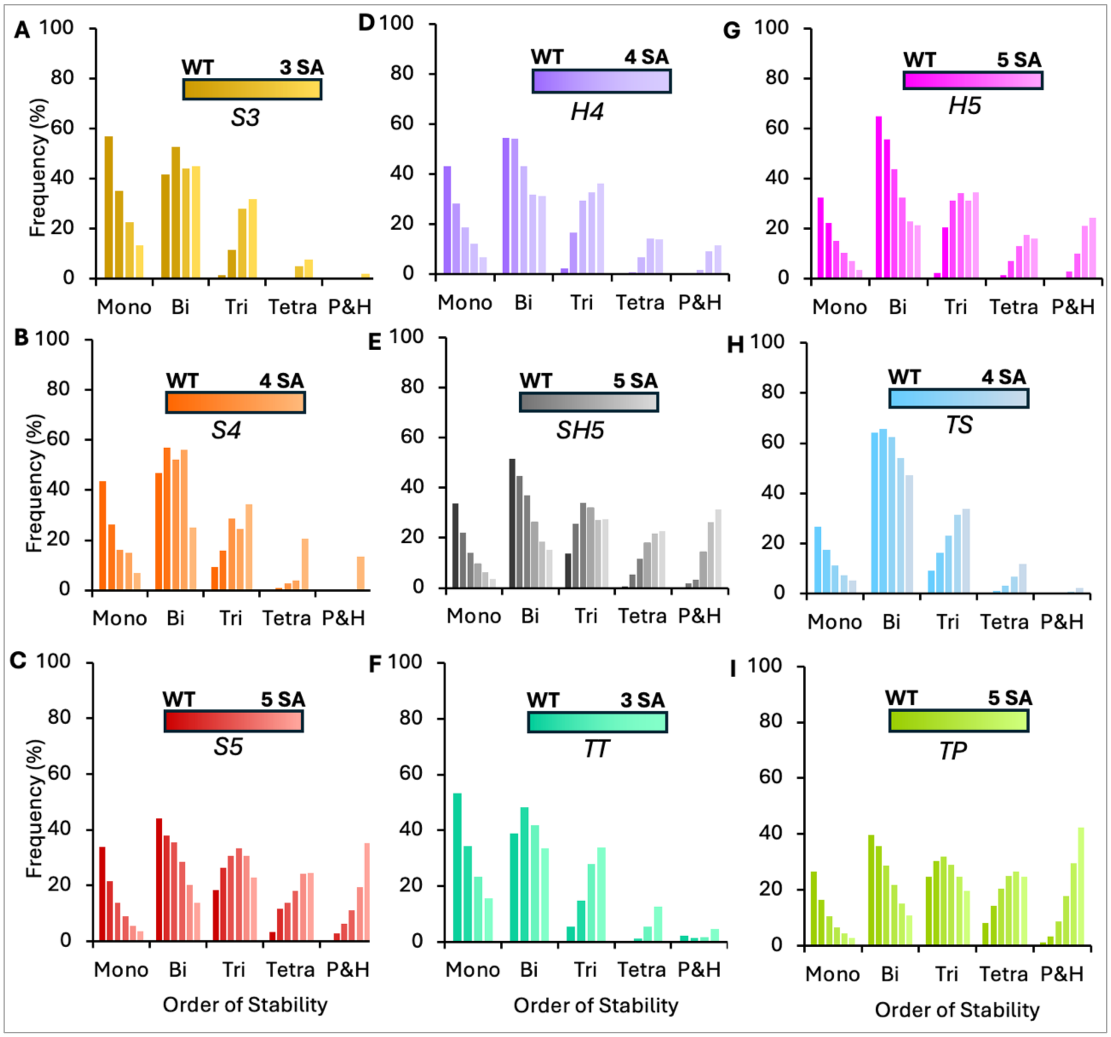

*Fig. S2 Bar plots showing the steady-state distribution (SSD) of all the WT networks and their self-activated counterparts. The first vertical panel contains HDFLs having serial topology, S3, S4, S5. As we move along the first horizontal panel, network becomes more hub-type. At the center, SH5 is a mixed serial/hub network. Also, TT, TS, TP are cyclic HDFLs with three, four, and five nodes. The left-extreme of the color gradient panel in each network represents WT (unperturbed) network. As the color gets darker, the number of self-activations in the network increases. The right-extreme of the color gradient panel represents the network with all nodes self-activated.*

### S3. Topologically distinct networks with the same number of nodes can exhibit different dynamics

The data reveals that topologically distinct networks with the same node count can exhibit different numbers of alternative states. It also shows that each cyclic network allows an extra cluster than the serial network with the same number of nodes. For instance, TT allows three while S3 allows two attractors (top horizontal panel), TS possibly allows four while S4 allows three attractors (middle horizontal panel), and TP exhibits five while S5 exhibits four attractors (lower horizontal panel). This shows that cyclic networks operate closer to serial networks and both of these operate opposite to hub-type networks (H4, H5) which show only a few attractors (just two).

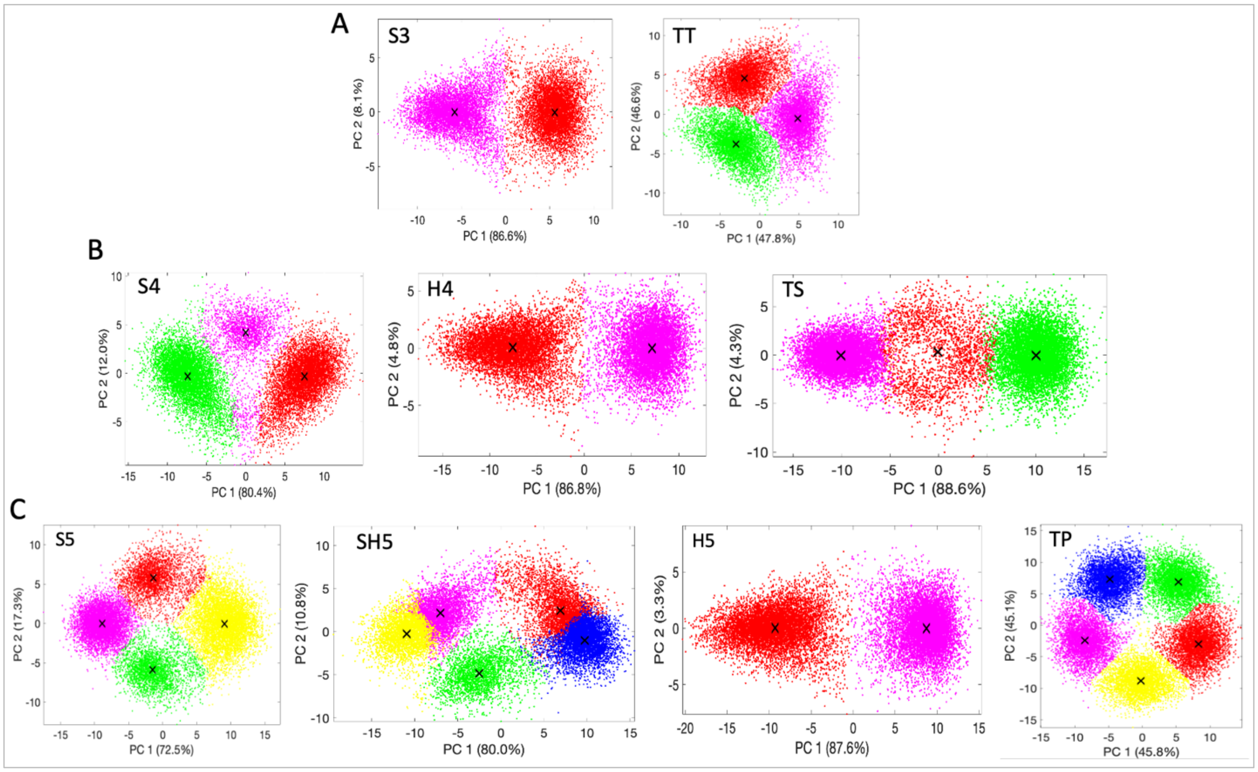


*Fig. S3 Comparative analysis of alternative states (clusters) in topologically distinct networks with the same number of nodes. (A) Three-node networks show two (in S3) and three (in TT) states. (B) Four-node networks show three (in*

*S4), two (in H4), and three to four (in TS) states. (C) Five-node networks four (in S5), five (in SH5), two (in H5), and five (in TP) states.*

### S4. Perturbations to restrict the dimension of attractor space of the networks.

Here we analyze the effect of single-edge perturbations (ED and ESR) on the SSD of HDFLs. We take an HDFL and randomly delete an edge or change the sign of an edge from repression to activation. We simulate the two perturbed networks and compare their SSD with the WT counterpart. A similar procedure is carried out throughout all HDFLs. For all networks, the data emphasize that both edge perturbations significantly increase lower-order stability which is compensated by a decrease in higher-order stability. The SSD peak therefore shifts to the left. We also observe that an ESR is more effective than an ED in increasing lower-order stability since each ESR results in more frequency of lower-order stability than an ED. Furthermore, the data depicts that when all the network nodes are self-activated single ED and single ESR become ineffective in reducing (increasing) the frequencies of higher-order (lower-order) attractors.

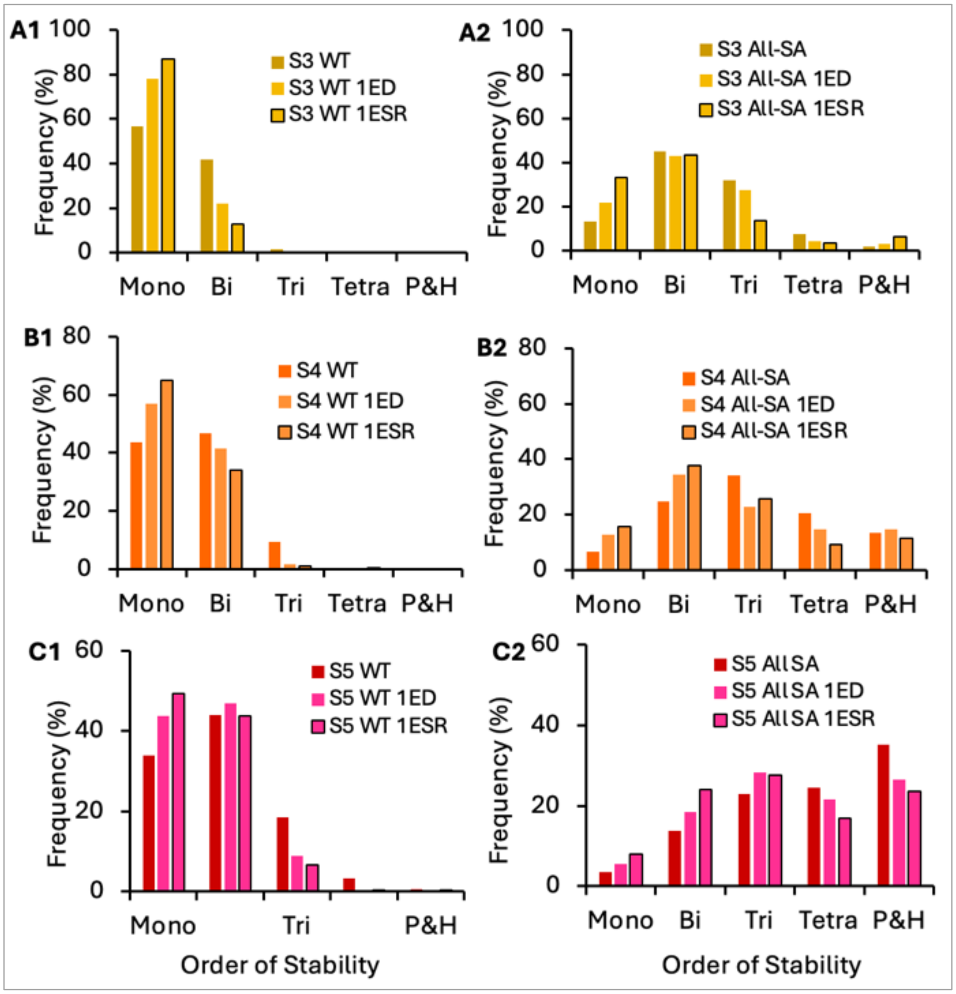


*Fig. S4 Effect of single Edge Deletion (ED) and single Edge Sign Reversal (ESR) on the SSD of Serial-Type HDFLs. (A1-C1) Bar graphs showing comparison of SSDs of WT Serial HDFLs with their two perturbed networks. (A2-C2) Same as left panel, except that all the network nodes are self-activated.*

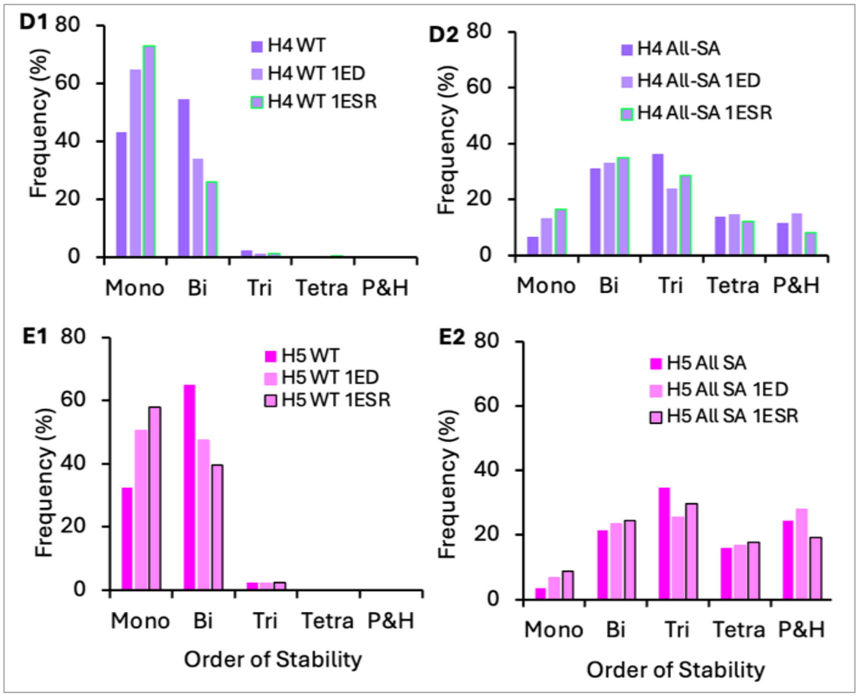


*Fig. S5 Effect of single Edge Deletion (ED) and single Edge Sign Reversal (ESR) on the SSD of Hub-Type HDFLs. (A1-C1) Bar graphs showing comparison of SSDs of WT Hub HDFLs with their two perturbed networks. (A2-C2) Same as left panel, except that all the network nodes are self-activated.*

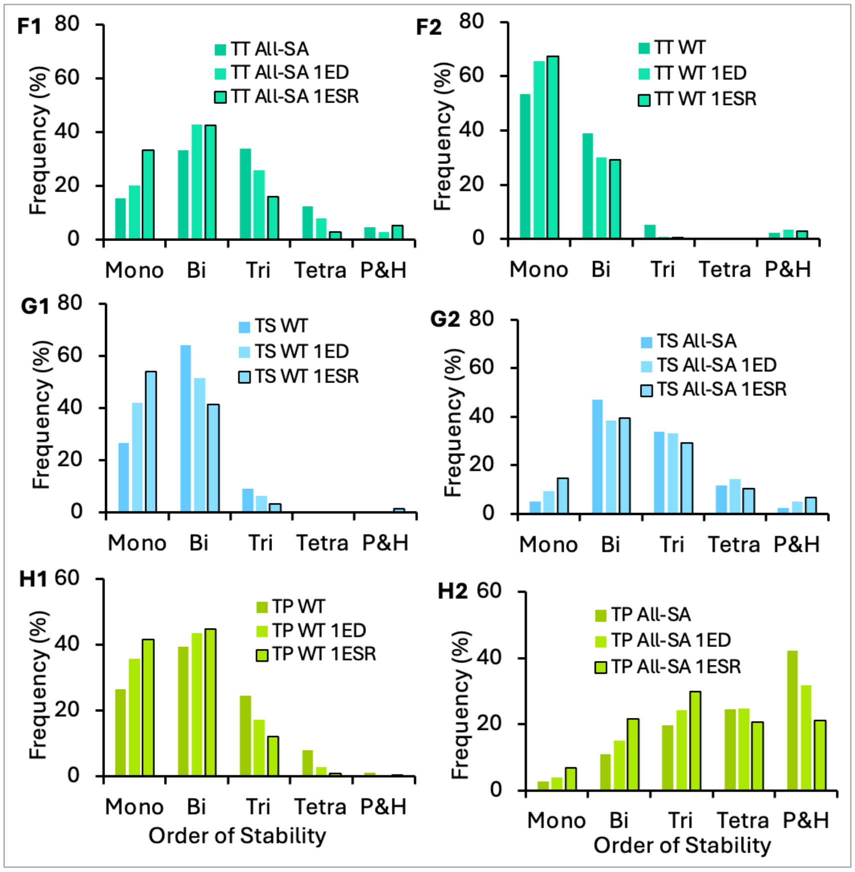


*Fig. S6. Effect of single Edge Deletion (ED) and single Edge Sign Reversal (ESR) on the SSD of Cyclic HDFLs. (A1-C1) Bar graphs showing comparison of SSDs of WT Cyclic HDFLs with their two perturbed networks. (A2-C2) Same as left panel, except that all the network nodes are self-activated.*

### S5. Effect of increasing the number of edge perturbations.

Here we identify the number of perturbations that are required to convert a multi-attractor (multistable) HDFL into a single-attractor (monostable) HDFL. In general, we see that we need to break all PFLs (equivalent to the number to toggle switches) in the network to achieve monostablity. Since serial-and hub type networks having $n$ nodes contain $(n-1)$ PFLs. In other words, in an $n$ node serial or hub type network, $(n-1)$ PFLs need to be eliminated to achieve unique steady state. Further, any cyclic network has as many nodes as toggle switches and we observe that even after deleting all PFLS these networks still don't become completely monostable, though the frequency of monostable attractors is the highest after breaking all the loops. This may be attributed to a unique property of cyclic networks as even after breaking all loops, there still remains the possibility of the existence of a "global" feedback (involving all nodes) or a feedforward loop in these networks. On the other hand, we see that in networks with all the self-activated nodes, only mono or bistable attractors is never possible. In fact, with maximum perturbations, HDFLs show almost equal possibilities for mono, bi, and tristable attractors. This again reiterates that networks with autoregulated nodes always tend to exhibit higher-order stability.

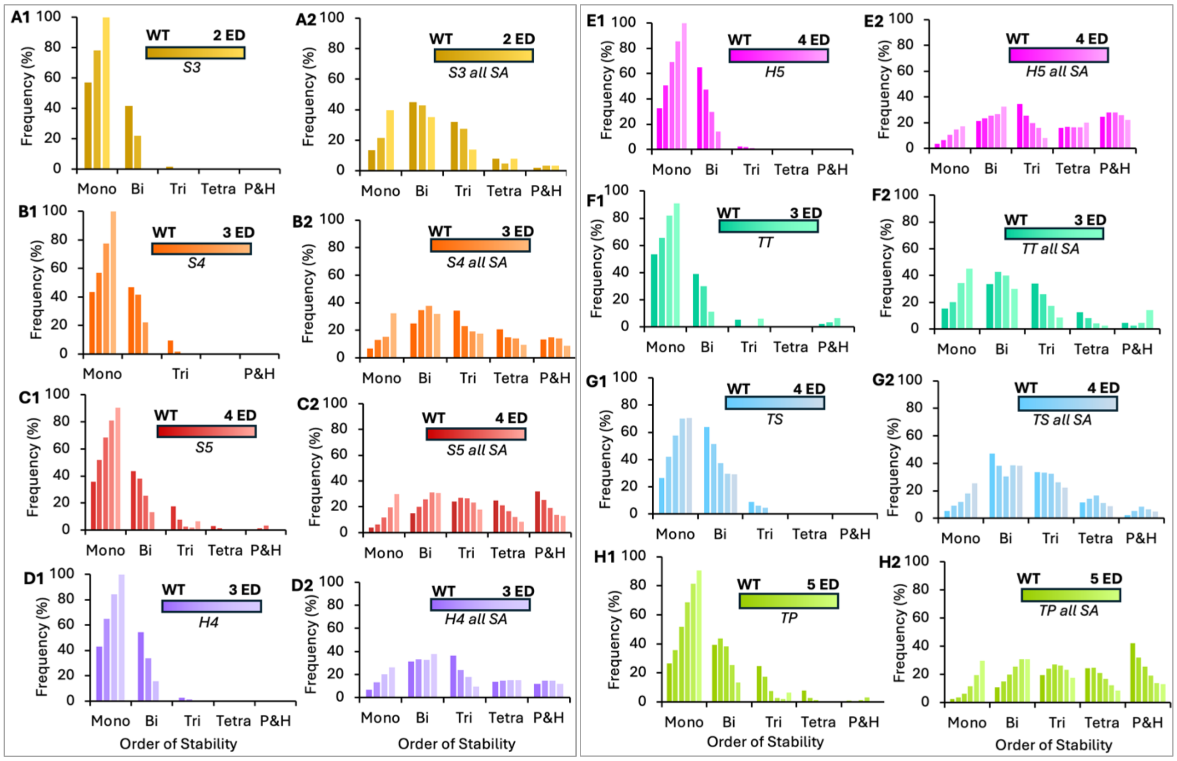


*Fig. S7 Comparision of the effect of multiple edge deletions on SSD in WT and self-activated HDFLs. (A1-H1) WT HDFLs undergo multiple edge deletions that result into increase in the frequency of lower-order states, and in some cases a fully monostable network. (B1-H1) Multiple edge deletions in the same HDFLs with all nodes self-activated create almost equal chances for mono-and bistable attractors in small-size networks and equal possibility for mono, bi, and tristable attractors in relatively large networks. "WT" in A2-H2 means "WT + all nodes self-activated".*

### S6. Effect of the position of edge EDs and ESRs on network dynamics.

Each of the eight networks in Fig. S7 has a self-activated (SA) terminal node and are referred to as asymmetric networks. An ED or ESR is applied at two positions in a network: P1 refers to an ED or ESR between two nodes involving a SA node. P2 refers to an ED or ESR at the other terminal of the network, i.e. the terminal that doesn't have a SA node. The data shows that an ED at P1 can increase (decrease) more monostable (bistable) states than an ED at P2. This effect becomes more pronounced when an ED is replaced by an ESR. The data thus reveals that the position of perturbation has functional significance in networks having SA nodes (asymmetric networks). It also highlights that gene interactions that involve autoregulated genes can serve as prime targets to control multiple alternative states during fate transition.

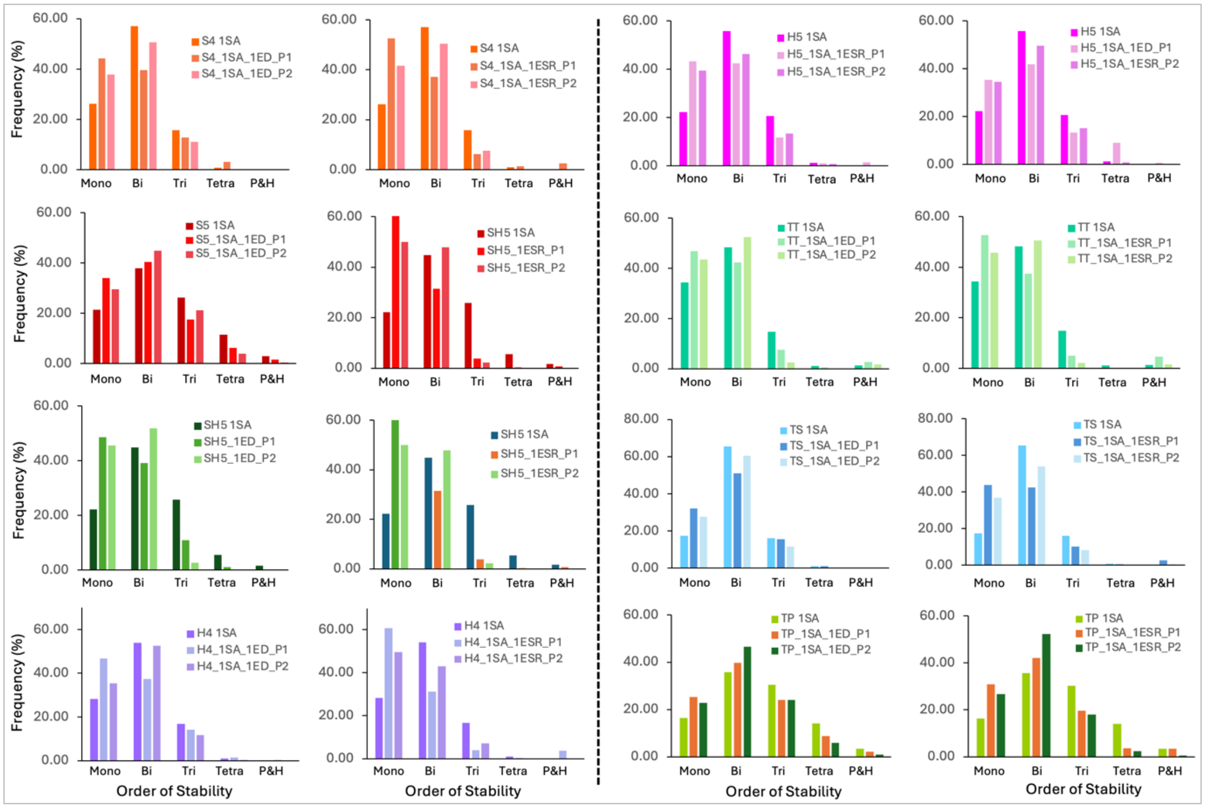


***Fig. S8 Bar plots showing the effect of the position of EDs and ESRs in asymmetric networks.*** *In each network, 1SA denotes one node is self-activated and that node is either of the terminal nodes. P1 and P2 denote the positions of ED or ESR. Each network is tested with ED and ESR at positions P1 and P2 and the SSD is compared with their unperturbed counterparts (i.e., network name_1SA). An ED or ESR at P1 reduces more bistable states and increases more monostable states than an ED or ESR at P2. Further, an ESR has a more pronounced effect than an ED. The left vertical panels show the effect of EDs at P1 and P2. The right vertical panels show the effect of ESRs at P1 and P2.*
